# The Prefrontal Cortex to the Hippocampus Glutamatergic Pathway Regulates Reward Effects of Methamphetamine

**DOI:** 10.1101/2024.05.20.595065

**Authors:** Dongdong Zhao, Wenjing Shi, Minyu Li, Rui Xue, Hang Wang, Le Zhang, Mengbing Huang, Liping Bai, Rou Gu, Ye Li, Xianwen Zhang, Peter Kalivas, Jie Bai

## Abstract

Methamphetamine (METH) induces reward effects. The prefrontal cortex (PFC) plays a role in regulating top-down. The hippocampus is closely involved reward memory. However, the direct projects from the PFC to the hippocampus have not been studied well. Here, we use conditioned place preference (CPP), a recombinant adeno-associated virus 2/9 (rAAV2/9) and optogenetic approaches to explore a role in METH-CPP for a novel pathway projecting from the medial prefrontal prelimbic cortex (PrL) directly to the dorsal hippocampus CA1 (dCA1). Expressing CaMKIIα-ChR2 or CaMKIIα-eNpHR3.0 in the PrL and optically stimulating in the dCA1 blocks or enhances METH-CPP, respectively. Moreover, silencing the PrL to dCA1 glutamatergic pathway with tetanus neurotoxin (TeNT) enhances METH-CPP. These results identify a novel PrL-to-dCA1 glutamatergic pathway regulating METH-CPP.

## Introduction

Methamphetamine (METH) is the one of most common illicit drugs worldwide, with 33 million users. Drug consumption is driven by drug rewarding effects. The process of reward to developing METH use disorder involves various brain regions^[1]^. The prefrontal cortex (PFC) is important for top-down regulation of motivated behaviors^[2]^, including reward seeking^[3]^. The PFC processes the value of a reward signal^[4]^, and PFC dysfunction has been implicated in the loss of control over drug use^[5]^. In rodents, the medial prefrontal cortex (mPFC) has been implicated in bidirectionally regulating reward seeking^[6]^. A common experimental measure of seeking is the capacity of addictive drugs to induce conditioned place preference (CPP), and activity in the dorsal hippocampus CA1 (dCA1) region has been implicated as critical in CPP^[7]^. Given the importance of both the PFC and dCA1 in reward seeking it is surprising that a direct pathway, the mPFC to the dCA1 has not been closely studied in regulating drug induced CPP. Here, we show that a subset of mPFC projections into to the dCA1 bidirectionally regulates METH induced CPP.

### Direct mPFC projection to the dCA1

To investigate the presence of a projection from the mPFC to the dCA1, we used brain stereotaxic surgery to inject a recombinant adeno-associated virus 2/9 (rAAV2/9)-human Synapsin I (hSyn)-mCherry, the anterograde non-transsynaptic tracer^[8]^ (**Figure 1a**) into the mPFC in mice (Figure 1b). Then the mice were placed back in the home cage for 3 weeks to allow for the virus expression to reach its maximum (Figure 1c). The fluorescent protein (mCherry, red) was expressed in the mPFC neurons (Figure 1d) and mCherry-containing terminals were expressed in the dCA1 (Figure 1e). Other the mPFC axon terminal regions were also observed, such as the olfactory bulb (onl), striatum (STRd) and others (**Supplemental Figure 1**). These results show a direct projection from the mPFC to the dCA1 in mice.

**Figure 1.**
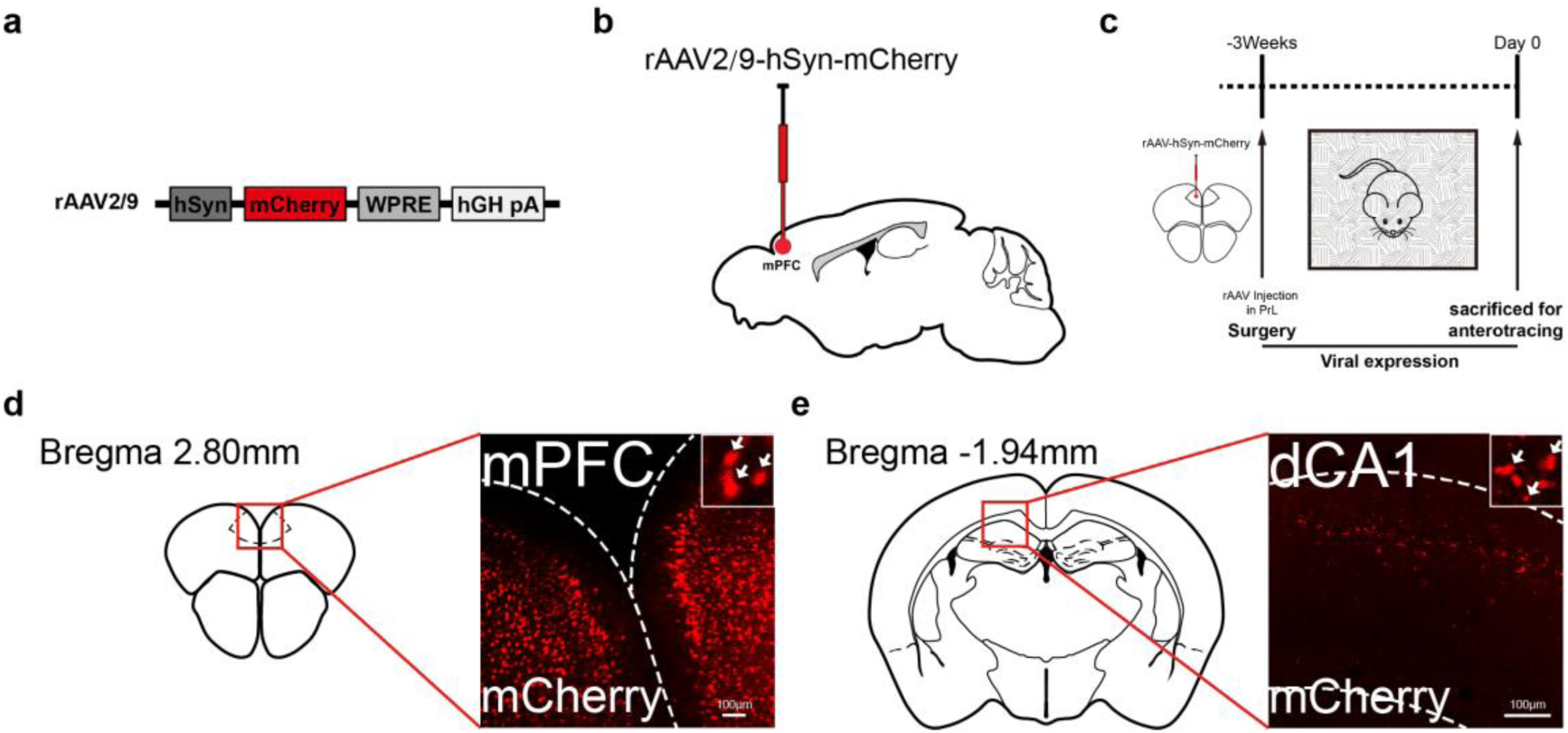
The mPFC to the dCA1 projection. **a**, Anterograde tracing rAAVs. **b**, Diagram of anterograde virus injection site in the mPFC. **c**, Schematic of the experimental timeline of virus expression. **d**, The mCherry (red) fluorescent expressions in the mPFC (n=3 mice). The white box represents a magnified view of a specific area, the white arrow indicates the neuronal cell body of the mPFC. **e**, Representative image of the mCherry fluorescent targeting in the dCA1. The white box represents a magnified view of a specific area, the white arrow indicates the terminals from the mPFC neurons. mPFC, medial prefrontal cortex, dCA1, dorsal hippocampus CA1. Scale bar, 100 μm.

### The PrL-to-dCA1 top-down regulates CPP

The mPFC is composed of multiple sub-compartments and the prelimbic (PrL) subregion is a critical brain region for drug reward-seeking^[9]^. Activity of dCA1 is necessary for drug-induced conditioned place preference (CPP)^[7b]^, a common and reliable behavioral assay to assess the rewarding properties of drugs. Since we found that the dCA1 received input from the PrL, we asked whether the PrL to the dCA1 was involved in CPP by selectively activating or inhibiting the PrL to the dCA1 pathway with optogenetic stimulation of channelrhodopsin-2 (ChR2), a blue-light sensitive ion channel mCherry fluorescent protein (ChR2-mCherry) or the silencing opsin halorhodopsin (eNpHR3.0-mCherry), a yellow-light sensitive ion channel. To investigate the PrL to the dCA1 pathway regulating the place preference, rAAV2/9s expressing distinct transgenes including control rAAV2/9-mCherry, rAAV2/9-hSyn-hChR2(H134R)-mCherry and rAAV2/9-hSyn-eNpHR3.0-mCherry (**Figure 2a**) were microinjected into the PrL, optical fibers were implanted into the dCA1 (Figure 2b, c). A schematic of experimental timeline for the rAAV2/9s expression and CPP regulated by optogenetics was shown in Figure 2d. Three weeks after surgery mice were entered CPP training. Optical stimulation was applied only when mice were put into one side of the CPP apparatus (light-paired) for 20 min, whereas the light was turned off when mice were put into the other side for every other day for 8 days. On the 12th day, mice were free to either side. Real time place preference (RTPP) tracking was used to detect mice preference when the light stimulation was given (Figure 2e). Mice undergoing ChR2 stimulation of the PrL to the dCA1 pathway spent less time in the light-paired side compared to control mice injected with a control rAAV2/9 (ChR2= −60.28 ±44.35s; Control= −114.91 ±39.22s, respectively) (Figure 2f) although there is not a significant difference. Conversely, mice undergoing the inhibitory eNpHR3.0 stimulation spent more time in the light paired side (eNpHR3.0= 100.37 ±44.65 s; Control= −114.91 ±39.22s, respectively) (Figure 2g). Expression of mCherry in the PrL and in the dCA1 was shown in Figure 2h, i. The results suggest that activation of the PrL to dCA1 direct neural pathway is involved in CPP behavior in mice.

**Figure 2.**
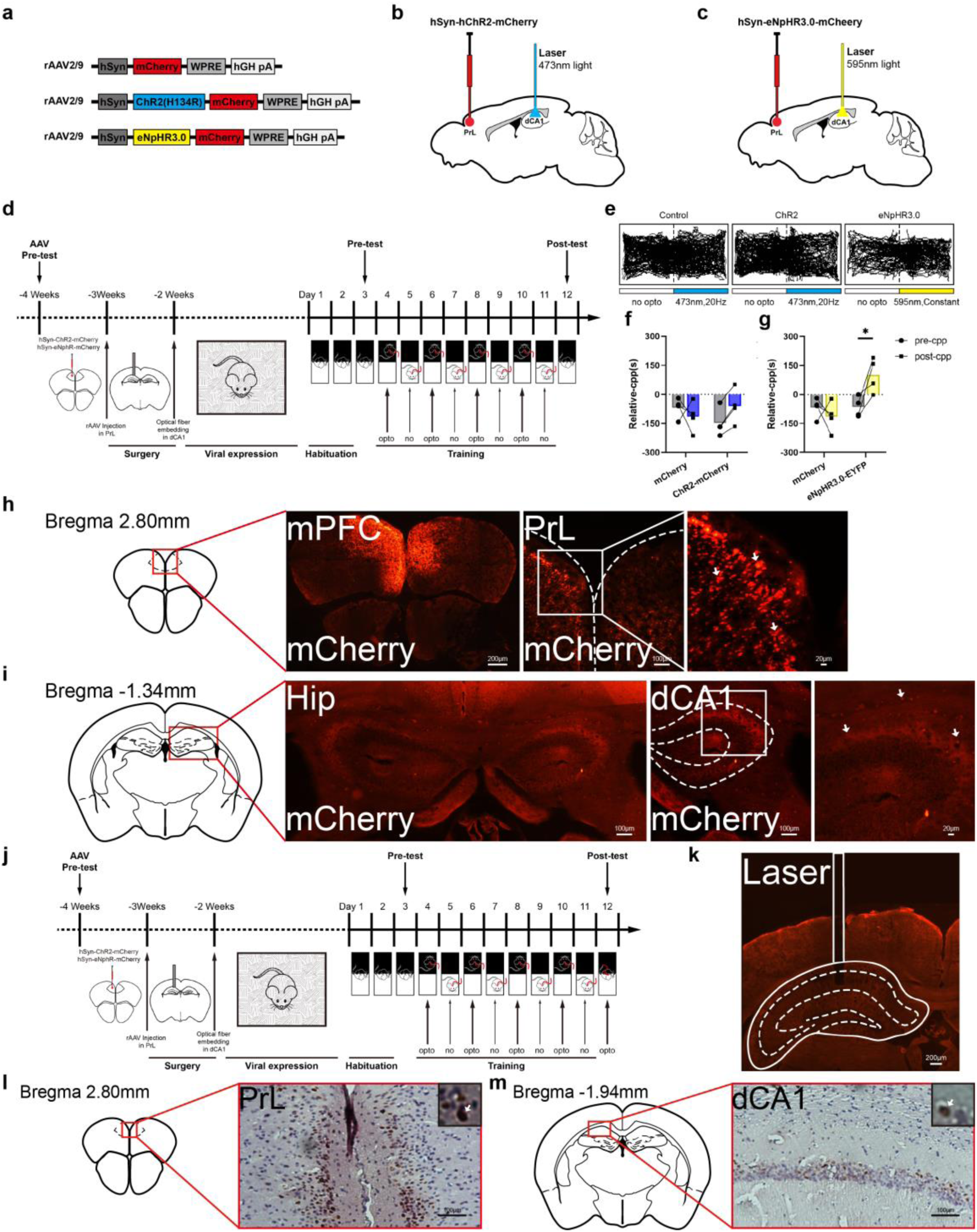
The CPP expression after the PrL to the dCA1 activation and inhibition. **a**, Anterograde tracing rAAVs. **b**, **c**, The schematic of the anterograde tracing strategy. Diagram of virus injection site in the PrL and optical fiber implantation site in the dCA1, 473 nm, blue light (**b**), 595 nm, yellow light (**c**). **d**, The schematic of the experimental timeline of rAAVs expression and CPP test combining with light stimulation. **e**, RTPP track of mice when the laser on. **f**, CPP expression in mCherry (n=4 mice) vs. ChR2-mCherry expressing mice (n=4 mice). (two-way analysis of variance (ANOVA), interaction, *F*_(1,12)_=3.231, *P*=0.0974; mCherry, *F*_(1,12)_=0.1089, *P*=0.7471; ChR2-mCherry, *F*_(1,12)_=0.3050, *P*=0.5909, Bonferroni’s multiple comparisons test). **g**, CPP expression in mCherry (n=4 mice) vs. eNpHR3.0-mCherry expressing mice (n=4 mice). (two-way ANOVA, interaction, *F*_(1,12)_=9.150, *P*=0.0106; mCherry, *F*_(1,12)_=9.693, *P*=0.0090; eNpHR3.0-mCherry, *F*_(1,12)_=2.902, *P*=0.1142, Bonferroni’s multiple comparisons test). **h**, **i**, Expression of mCherry in the PrL (**h**) and dCA1 (**i**) (n=3 mice). PrL, prelimbic of medial prefrontal cortex. Scale bar, 100μm, 20μm. **j**, The schematic of the experimental timeline of rAAVs expression and CPP test combining with light stimulation on 12th day (with opto). **k**, Diagram of laser fiber embedding site on dCA1. **l**, **m**, The c-fos expression in the PrL and dCA1 with ChR2 blue stimulation (n=3 mice). Scale bar, 100 μm. Data in **f**, **g** are presented as means±s.e.m. Two-sided statistical tests were used. *\*p* < 0.05.

To examine functional validation of the ChR2, a schematic experimental timeline for the rAAV2/9s expression and CPP regulated by optogenetics in an additional cohort of mice was shown in Figure 2j and the site of laser fiber implantation in the dCA1 shown in Figure 2k. 90 min following optical stimulation (473 nm, 3.0 mW) on 12th day the expression of c-Fos, a marker of neuronal activity^[10]^, was detected in the PrL and dCA1 of mice. We could see that the expression of c-Fos in the PrL and dCA1 was detected (Figure 2l, m). The results suggest that validation of the ChR2 in the PrL-to-dCA1 neural pathway induces neuronal activity.

These results indicate that directly inhibiting the PrL to the dCA1 pathway is rewarding and produces CPP, while activating the pathway reduced CPP. This suggests possibility that tonic activity in the PrL-to-dCA1 pathway may negatively regulate the development of CPP.

### The PrL to the dCA1 optogenetic regulation on METH-CPP

The experiment in Figure 2 shows that activity in the PrL-to-dCA1 neural pathway may negatively regulate CPP in mice. To test whether regulating activity in the PrL to the dCA1 pathway would alter METH-CPP, we employed the same rAAV2/9s (**Figure 3a)** and CPP protocol used in Figure 2, except that mice were injected with METH (2.5 mg/kg) or saline on alternate days in distinct sides of the apparatus. For the experiments mice were habituated for 2 days before obtaining a Pre-CPP measurement (Figure 3d). In the first experiment with METH-CPP, rAAV2/9s injection in the PrL and projection to the dCA1 is shown in Figure 3b for ChR2 and 3c for eNpHR3.0. METH induced CPP in mice receiving either transgene when no optical stimulation was applied. However, when optical stimulation was applied again 24 hours later, stimulation of ChR2 in the PrL projects to the dCA1 reduced CPP (Figure 3e), while inhibition of eNpHR3.0 in the PrL projects to the dCA1 potentiated CPP (Figure 3f) (ChR2= −119.58 ± 52.08 s; eNpHR3.0= 204.37 ± 50.67 s; Control= 46.80 ±36.55 s, respectively). Note the overlap of mCherry with neuron soma-specific DAPI in the PrL injection site and the lack of overlaps between the mCherry terminals and DAPI in the dCA1 (Figure 3g, h). In either experiment METH-CPP was unaffected by stimulation in control mice transfected with control rAAV2/9. Since the first experiment shown in Figure 3e, f possibly contains an ordering effect that stimulation was always given after no optical stimulation on 12th day during the test for METH-CPP on 13th day, an additional experiment was conducted to where only optical stimulation was administered during the test for CPP on 12th day. The experimental schematic for the rAAV2/9s and METH-CPP procedure regulated by optogenetics was shown (Figure 3i). As in the first experiment, METH-CPP was induced in control mice, and this was blocked by optical stimulation in the ChR2 mice and potentiated in the eNpHR3.0 mice (Figure 3j, k). (ChR2= −84.39 ±29.41 s; eNpHR3.0= 256.14 ±32.64 s; Control= 117.98 ±42.50 s, respectively). Figure 3l, m shows anatomical validation of rAAV2/9s injection in the mPFC and terminal mCherry in the dCA1. These results indicate that directly activating the PrL to the dCA1 pathway inhibits METH-CPP, while inhibiting the pathway enhances METH-CPP. This suggests possibility that activity in the PrL-dCA1 pathway may negatively regulate the development of METH-CPP.

**Figure 3.**
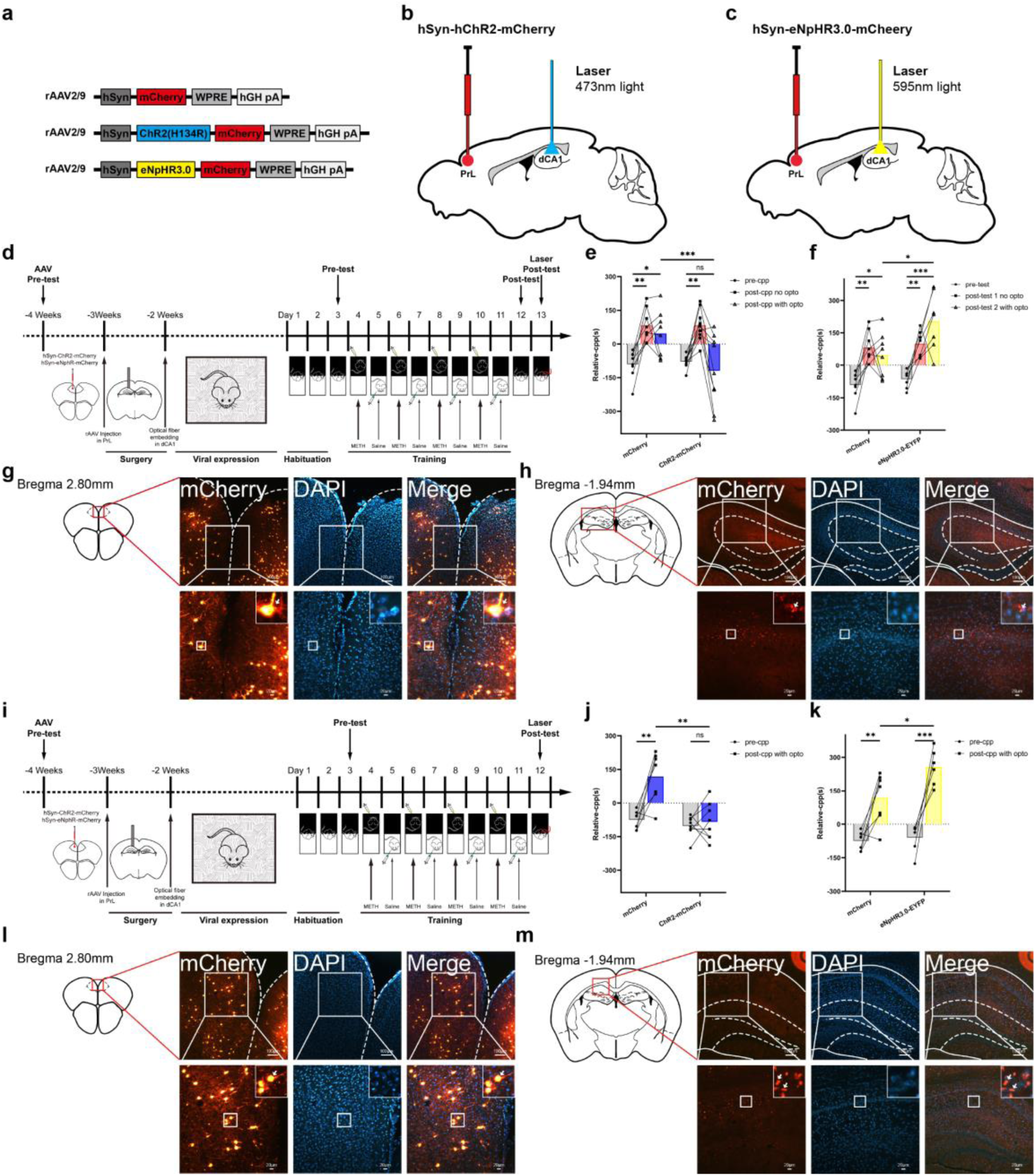
The PrL- dCA1 optogenetic regulation on METH-CPP. **a**, Anterograde tracing rAAVs. **b**, **c**, The schematic of the anterograde tracing strategy. Diagram of virus injection site in the PrL and optical fiber implantation site in the dCA1. **d**, The schematic of the experimental timeline of rAAVs expression and METH-CPP test on 12th day (no opto) combining with light stimulation on 13th day (with opto). **e**, METH-CPP expression in mCherry (n=8 mice) vs. ChR2-mCherry expressing mice (n=9 mice). (two-way ANOVA, interaction, *F*_(2,45)_=4.958, *P*=0.0113; CPP (pre-, post-no, post-with opto), *F*_(1,45)_=3.955, *P*=0.0528; mCherry (n.c, ChR2), *F*_(2,45)_=14.95, *P*<0.0001, Bonferroni’s multiple comparisons test), n.c, negative control group. **f**, METH-CPP expression in mCherry (n=8 mice) vs. eNpHR3.0-mCherry expressing mice (n=7 mice). (two-way ANOVA, interaction, *F*_(2,39)_=3.210, *P*=0.0512; CPP (pre-, post-no, post-with opto), *F*_(1,39)_=7.004, *P=*0.0117; mCherry (n.c, eNpHR3.0), *F*_(2,39)_=24.71, *P=*0.0001, Bonferroni’s multiple comparisons test), n.c, negative control group. **g**, **h**, Expression of mCherry and DAPI in the PrL (g) and dCA1 (h) (n = 3 mice), DAPI, 4′,6-diamidino-2-phenylindole, Scale bar, 100 μm, 20 μm. **i**, The schematic of the experimental timeline of rAAVs expression and METH-CPP test on 12th day (with opto). **j**, METH-CPP expression in mCherry (n=7 mice) vs. ChR2-mCherry expressing mice (n=8 mice). (two-way ANOVA, interaction, *F*_(1,26)_=9.920, *P*=0.0041; CPP (pre-, post-with opto), *F*_(1,26)_=17.21, *P*=0.0003; mCherry (n.c, ChR2), *F*_(1,26)_=14.61, *P*=0.0007, Bonferroni’s multiple comparisons test), n.c, negative control group. **k**, METH-CPP expression in mCherry (n=7 mice) vs. eNpHR3.0-mCherry expressing mice (n=6 mice). (two-way ANOVA, interaction, *F*_(1,22)_=3.902, *P*=0.0609; CPP (pre-, post-with opto), *F*_(1,22)_=5.981, *P*=0.0229; mCherry (n.c, eNpHR3.0), *F*_(1,22)_=66.58, *P* < 0.0001, Bonferroni’s multiple comparisons test), n.c, negative control group. **l**, **m**, The expression of mCherry and DAPI in the PrL (**l**) and dCA1 (**m**) (n = 3 mice). Scale bar, 100 μm, 20 μm. Data in **e**, **f**, **j**, **k** are presented as means±s.e.m. Two-sided statistical tests were used. *\*p*<0.05, *\*\*p*<0.01, *\*\*\*p*<0.001; ns, not significant.

### Optogenetic activation or inhibition of glutamatergic PrL-to-dCA1 reduces or enhances METH-CPP

Although a significant body of evidence supports a role for PFC dopamine signaling pathway in drug-related rewarding. The specific circuitry that is associated with drug reward has been broadened to include many neural inputs and outputs that interact with the basal forebrain, such as glutamatergic projections from the mPFC^[11]^.

Hypofrontality in METH addiction may also relate to perturbations in PFC excitatory glutamate transmission^[12]^. We used brain stereotaxic surgery combined with injecting of rAAV2/9- Ca^2+^/calmodulin-dependent protein kinase II α (CaMKⅡα)-DIO-mCherry into the PrL, rAAV2/Retro-hSyn-Cre-EYFP into the dCA1 in mice (**Figure 4a, b**). Other areas from glutamatergic PrL projection and from the dCA1 input. Other the mPFC axon terminal regions were also observed, such as onl, STRd (**Supplemental Figure 2**). These results show a direct glutamatergic projection from the mPFC to the dCA1 in mice. The mice were placed back in the home cage for 3 weeks to optimize rAAV2/9s expression (Figure 4c). The mice were sacrificed, and brain tissues were taken to validate viral expression. The CaMKⅡα promoter was used to restrict expression largely in glutamatergic neurons, and fluorescent protein expression was observed in the PrL and dCA1. The mCherry and EYFP are co-expressed in the PrL (Figure 4d) and dCA1 (Figure 4e). The mCherry indicates projection from the PrL to the dCA1, whereas the EYFP indicates projection from the dCA1 to the PrL. The results identified a novelty PrL-to-dCA1 glutamatergic projection, which is different from previous study, in which long-range GABAergic projections from the PFC to the dorsal hippocampus^[13]^.

**Figure 4.**
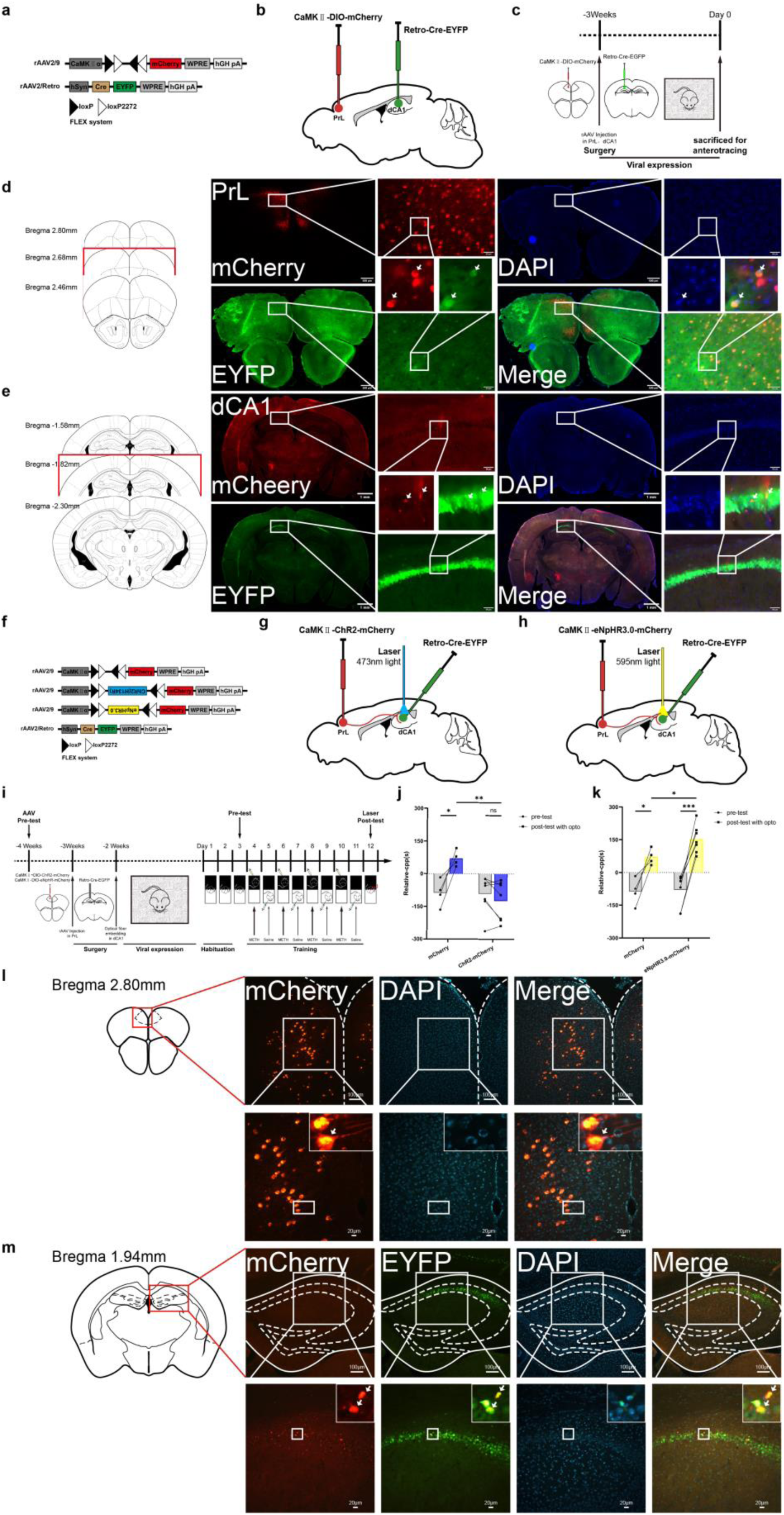
The glutamatergic PrL-to-dCA1 regulation on METH-CPP. **a**, Anterograde tracing rAAVs. **b**, The schematic of the DIO-Cre tracing strategy. **c**, Schematic of the experimental timeline of virus expression. **d**, **e**, The mCherry, EYFP and DAPI expression in the PrL (**d**) and dCA1 (**e**) (n= 3mice). Scale bar, 1mm, 500 μm, 50 μm. **f**, Anterograde tracing rAAVs. **g**, **h**, The schematic of the anterograde tracing strategy. Diagram of virus injection site in the PrL and optical fiber implantation site in the dCA1. **i**, The schematic of the experimental timeline of rAAVs expression and METH-CPP test combining with light stimulation on 12th day. **j**, METH-CPP expression in mCherry (n=4 mice) vs. ChR2-mCherry expressing mice (n=7 mice). (two-way ANOVA, interaction, *F*_(1,18)_=7.643, *P*=0.0128; CPP (pre-, post-with opto), *F*_(1,18)_=8.332, *P*=0.0098; mCherry (n.c, ChR2), *F*_(1,18)_=14.61, *P*=0.0837, Bonferroni’s multiple comparisons test), n.c, negative control group. **k**, METH-CPP expression in mCherry (n=4 mice) vs. eNpHR3.0-mCherry expressing mice (n=7 mice). (two-way ANOVA, interaction, *F*_(1,18)_=1.715, *P*=0.2068; CPP (pre-, post-with opto), *F*_(1,18)_=2.410, *P*=0.1380; mCherry (n.c, eNpHR3.0), *F*_(1,18)_=47.73, *P* < 0.0001, Bonferroni’s multiple comparisons test). **l**, **m**, The mCherry, EYFP and DAPI expression in the PrL (**l**) and dCA1 (**m**) (n=3 mice). Scale bar, 100 μm, 20 μm. Data in **j**, **k** are presented as means±s.e.m. Two-sided statistical tests were used. *\*p*<0.05, *\*\*p*<0.01, *\*\*\*p*<0.001; ns, not significant, n.c, negative control group.

Prefrontal functions are supported by local microcircuits involving long-range glutamatergic pyramidal neurons^[14]^ and prefrontal glutamate is involved in METH sensitization and preference^[15]^. Thus, we further examined whether enhancing or silencing glutamatergic activity in the PrL to the dCA1 pathway could block or enhance METH rewarding effect. The rAAV2/9s were injected into the PrL and dCA1 by brain stereotaxic surgery (Figure 4f), the sites of rAAV2/9-CaMKⅡα-DIO-hChR2(H134R)-mCherry, rAAV2/9-CaMKⅡα-DIO-eNpHR3.0-mCherry, and rAAV2/Retro-hSyn-Cre-EYFP with optical fiber (473 nm, 3.0 mW) are shown in Figure 4g, h. Experimental schematic for the rAAV2/9 and CPP procedure regulated by optogenetics were shown (Figure 4i). We detected the CPP after rAAV2/9s expression and METH (2.5 mg/kg) administration. METH-CPP was expressed in mice receiving control rAAV2/9-CaMKⅡα-DIO-mCherry, which was suppressed in mice receiving optogenetic activation of rAAV2/9-CaMKⅡα-DIO-hChR2(H134R)-mCherry (Figure 4j), but enhanced in mice receiving optogenetic inhibition of rAAV2/9-CaMKⅡα-DIO-eNpHR3.0-mCherry (Figure 4k) (ChR2= −124.81 ±34.56 s; eNpHR3.0= 151.93±24.37 s; Control= 70.73 ±18.20 s, respectively). The validation of mCherry in the PrL and the projection into the dCA1 is shown in Figure 4l, m. The results show that the direct glutamatergic projection from the mPFC to the dCA1 is novelty, which bidirectionally regulates METH-CPP in mice.

### Silencing the PrL-to-dCA1 glutamatergic pathway enhances CPP of METH

The PrL inactivation enhances cocaine seeking in certain circumstances^[16]^. Inactivation of the PrL cortex decreases discriminative stimulus-controlled relapse to cocaine seeking in rats^[17]^. To study the necessity of the PrL-to-dCA1 glutamatergic pathway in METH rewarding, the effect of knocking down the PrL to the dCA1 glutamatergic pathway on METH rewarding was further investigated by using tetanus toxin (TeNT) to impede release of glutamatergic synaptic vesicles. The rAAV2/9s containing rAAV2/9-CaMKⅡα-mCherry (**Figure 5a**) were injected into the PrL by brain stereotaxic surgery, rAAV2/Retro-hSyn-Cre-EYFP to the dCA1 (Figure 5b).

**Figure 5.**
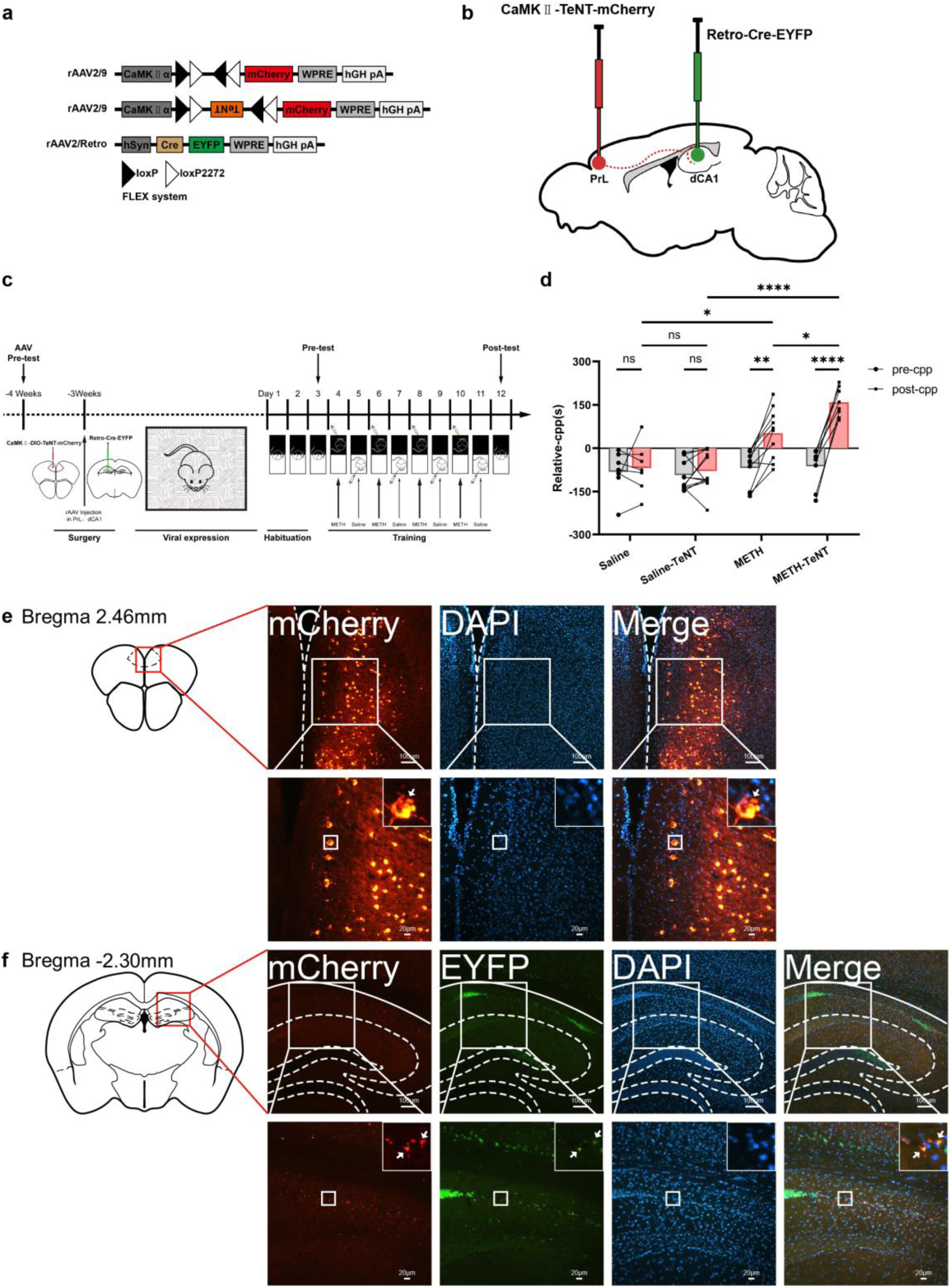
Glutamatergic PrL to dCA1 silencing enhances METH-CPP. **a**, Anterograde tracing rAAVs. **b**, Diagram of DIO-Cre virus injection site in the PrL and dCA1. Schematic showing the viral strategy to silence synaptic outputs of PrL to dCA1 neurons with tetanus toxin (TeNT). **c**, The schematic of the experimental timeline of rAAVs expression and METH-CPP test. **d**, METH-CPP expression of vehicle (n=10 mice) vs. TeNT (n=8 mice) expressing mice in Saline (n=19 mice) vs. METH groups (n=18 mice). (two-way ANOVA, interaction, *F*_(3,66)_=9.006, *P* < 0.0001; CPP (Saline, METH), *F*_(3,66)_=14.67, *P* < 0.0001; treated (n.c, TENT), *F*_(1,66)_=31.75, *P* < 0.0001, Bonferroni’s multiple comparisons test), n.c, negative control group. **e**,**f**, The mCherry, EYFP and DAPI expression in the in the PrL (**e**) and in the dCA1 (**f**) (n=3 mice). Scale bar, 100 μm, 20 μm. Data in **d** are presented as means±s.e.m. Two-sided statistical tests were used. *\*p*<0.05, *\*\*p*<0.01, *\*\*\*\*p*<0.0001; ns, not significant.

Experimental schematic for rAAV2/9s and METH-CPP procedure were as described in Figure 5c. Post-CPP was tested on the 12th day, METH-CPP was expressed in control mice injected with rAAV2/9-CaMKⅡα-DIO-mCherry, but was enhanced in mice expressing TeNT (Figure 5d) (Control= 51.66 ±27.58 s; TeNT= 158.31 ±17.71 s, respectively). Confocal images show expression of mCherry in the PrL (Figure 5e) and the projection to the dCA1 (Figure 5f). The results show that silencing PrL-to-dCA1 glutamatergic transmission enhances METH-CPP.

### The expression of glutamatergic receptor in silencing PrL-to-dCA1 glutamatergic pathway

To further determine the molecules involved in enhanced METH-CPP by reducing PrL-to-dCA1 activity, N-methyl-D-aspartate receptor 2B subunit (NMDAR2B), α-amino-3-hydroxy-5-methyl-4-isoxazole-propionic acid receptor (AMPAR), and the vesicular glutamate transporter1 (VGLUT1) expression was quantified in the mPFC and dCA1 by using western blotting (**Figure 6a-h**).

**Figure 6.**
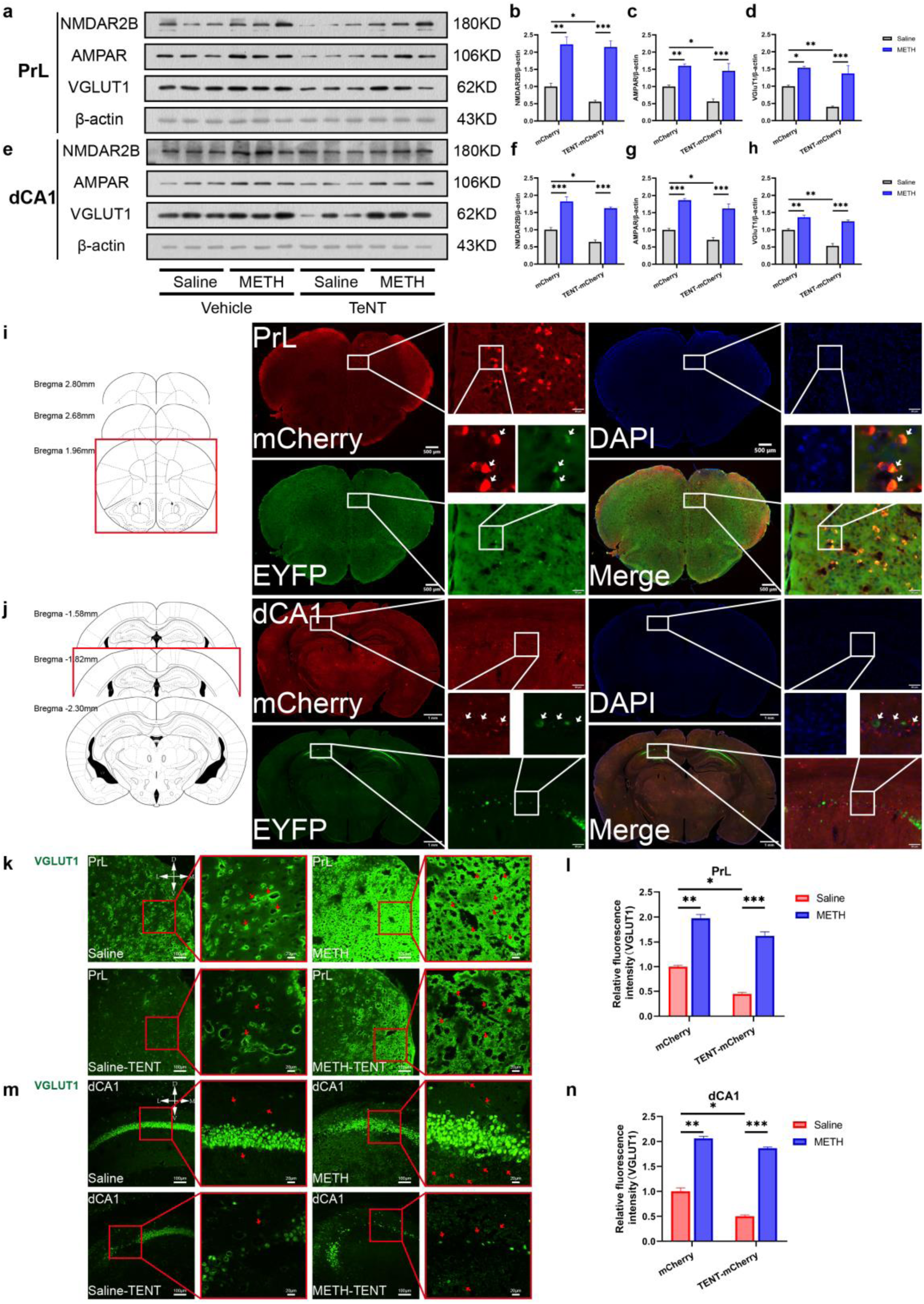
The expression of NMDAR2B, AMPAR and VGLUT1 in the PrL and dCA1. **a**, The expression of NMDAR2B, AMPAR and VGLUT1 in the PrL by western blotting. **b-d**, The relative expression on NMDAR2B, AMPAR and VGLUT1 in the PrL. **b**, shown by two-way ANOVA, interaction, F(1,18)=1.415, P=0.2497; CPP (Saline, METH), F(1,18)=2.869, P=0.1075; treated (n.c, TENT), F(1,18)=84.61, P < 0.0001, Bonferroni’s multiple comparisons test; **c**, shown by two-way ANOVA, interaction, *F*_(1,19)_=1.864, *P*=0.1881; CPP (Saline, METH), *F*_(1,19)_=7.467, *P*=0.0132; treated (n.c, TENT), *F*_(1,19)_=49.68, *P* < 0.0001, Bonferroni’s multiple comparisons test; **d**, shown by two-way ANOVA, interaction, *F*_(1,18)_=5.709, *P*=0.0280; CPP (Saline, METH), *F*_(1,18)_=18.71, *P*=0.0004; treated (n.c, TENT), *F*_(1,18)_=71.06, *P* < 0.0001, Bonferroni’s multiple comparisons test. **e**, The expression of NMDAR2B, AMPAR and VGLUT1 in the dCA1 by western blotting. **f-h**, The relative expression on NMDAR2B, AMPAR and VGLUT1 in the dCA1. **f**, shown by two-way ANOVA, interaction, *F*_(1,20)_=0.9132, *P*=0.3507; CPP (Saline, METH), *F*_(1,20)_=11.33, *P*=0.0031; treated (n.c, TENT), *F*_(1,18)_=119.9, *P* < 0.0001, Bonferroni’s multiple comparisons test; **g**, shown by two-way ANOVA, interaction, *F*_(1,20)_=0.0967, *P*=0.7590; CPP (Saline, METH), *F*_(1,20)_=10.45, *P*=0.0042; treated (n.c, TENT), *F*_(1,20)_=113.7, *P* < 0.0001, Bonferroni’s multiple comparisons test; **h**, shown by two-way ANOVA, interaction, *F*_(1,20)_=10.57, *P*=0.0040; CPP (Saline, METH), *F*_(1,20)_=29.16, *P* < 0.0001; treated (n.c, TENT), *F*_(1,20)_=99.83, *P* < 0.0001, Bonferroni’s multiple comparisons test; n.c, negative control group. **i**, **j**, Expression of mCherry, EYFP and DAPI in the PrL (**i**) and dCA1 (**j**) (n =3). Scale bar, 1mm, 100 μm, 50 μm. **k**, Confirmation by VGLUT1-immunofluorescence (Green) in the PrL. Scale bar, 100 μm, 20 μm. **l**, VGLUT1 fluorescent expression in the PrL by two-way ANOVA, interaction, *F*_(1,8)_=3.010, *P*=0.1210; CPP (Saline, METH), *F*_(1,8)_=60.36, *P* < 0.0001; treated (n.c, TENT), *F*_(1,8)_=341.5, *P* < 0.0001, Bonferroni’s multiple comparisons test). **m**, Confirmation by VGLUT1-immunofluorescence (Green) in the dCA1. Scale bar, 100 μm, 20 μm. **n**, VGLUT1 fluorescent expression in the dCA1 by two-way ANOVA, interaction, *F*_(1,8)_=11.76, *P*=0.0090; CPP (Saline, METH), *F*_(1,8)_=62.24, *P* < 0.0001; treated (n.c, TENT), *F*_(1,8)_=752.9, *P* < 0.0001, Bonferroni’s multiple comparisons test). Data in **b**, **c**, **d**, **f**, **g**, **h**, **l**, **n** are presented as means±s.e.m. Two-sided statistical tests were used. *\*p*<0.05, *\*\*p*<0.01, *\*\*\*\*p*<0.0001.

The expression of all three proteins in both PrL and dCA1 was increased by METH in control mice. The levels of the three proteins were reduced in both brain regions in TeNT mice treated with saline. Although control level of proteins was reduced by expressing TeNT in the PrL to the dCA1 pathway, the level of proteins after METH administration remained the same between TeNT and mCherry control mice, indicating that proportionally greater increase in proteins expression was produced by METH compared to saline treatment in TeNT mice. The expression of mCherry and EYFP in the PrL and dCA1 are shown in Figure 6i, j. VGLUT1 fluorescence staining in the PrL (Figure 6k, l) and dCA1 (Figure 6m, n) was shown. These data parallel the findings described above obtained by western blotting for VGLUT1. These results suggest that VGLUT1 is increased by METH in TeNT group more than in vehicle group.

NMDAR2B has an important role in the development of CPP of psychostimulant abuse. METH increase in NMDA receptor currents and an increase in NMDAR2B surface expression^[18]^. METH causes enduring changes within the mPFC including augmented burst firing within glutamatergic neurons^[19]^. Changes in extracellular glutamate concentration increase NMDA receptor expression, and adaptations in pre- and post-synaptic glutamate transmission^[15, 18, 20]^. METH exposure was associated with glutamate receptor and glutamate vesicular proteins^[21]^. The proteins responsible for glutamate uptake by vesicles are the vesicular glutamate transporter (VGLUT). It has been shown that VGLUT1 is dynamically regulated via a polysynaptic pathway to facilitate vesicular accumulation of glutamate for subsequent release after METH^[22]^. VGLUT1 represents the best marker for corticostriatal glutamatergic neurons^[23]^. Thus, our results indicate that increase in glutamatergic molecules in METH treated TeNT group is related to enhanced METH-CPP after silencing the PrL- to-dCA1 pathway.

## Discussion

Here we define a function of new top-down glutamatergic pathway from the PrL region of the mPFC directly to the dCA1 in the control of METH rewarding behaviour. The pathway from the PrL to the dCA1 is capable of bidirectionally regulating the expression of METH-CPP. Thus, expressing ChR2 or eNpHR3.0 in the PrL and optogenetically stimulating or inhibiting the pathway blocked or enhanced METH-CPP, respectively (Figure 4j, k). Moreover, silencing of the PrL-to-dCA1 glutamatergic pathway with TeNT enhanced the METH-CPP (Figure 5d). This pathway is novelty and distinct from the well-studied reciprocal pathway of from the hippocampal-to-prefrontal pathway.

The mPFC is an important brain region that interacts with several brain areas^[24]^. Many of the pathologies driven by METH use involve cortical neural plasticity to impacting both top-down and bottom-up processing^[25]^ and restoring prefrontal cortex hypoactivity prevents compulsive addiction seeking^[16a]^. Recent work shows that discrete neural ensembles within the PrL are involved in the expression and inhibition of conditioned responding for drug rewards^[9a, 26]^. Also, METH administration induces changes in the object recognition memory circuit^[27]^ and dysregulated glutamate in the mPFC. Our results are generally consistent with the literature and more selectively show that at least for METH place conditioning the glutamatergic projection from the PrL to the dCA1 provides a critical regulatory role in how mice respond to conditioned environments. These findings have important implications for understanding the basic neurobiology of rewarding processing.

Although we are the first to demonstrate direct involvement of a glutamatergic projection from the PrL to the dCA1 in conditioning to addictive drugs, anatomical connections between the PFC and the hippocampus are known^[28]^. The mPFC influence on the CA1 activity predicts subsequent learning speed^[29]^ and oscillatory stimulation of the ventral hippocampus and mPFC facilitates neural transmission in the hippocampal-prefrontal pathway^[30]^. Moreover, the hippocampus, prefrontal cortex, and interconnected neural pathways are implicated in several aspects of cognitive and memory processes^[28b]^. Prelimbic neurons contribute to adjusting decisions based on internal valuations^[31]^. Pyramidal neurons selectively fire when animals reach certain positions in an environment in the CA1 of mice while the animals performed a goal-directed navigation task^[32]^. The hippocampus forms spatial codes that are critical for navigation, which are based on features of the environment, including sensory cues and the location of rewards^[33]^. Together, the literature supports our discovery that the PrL-to-dCA1 glutamatergic pathway regulates the conditioned location of rewards.

Although the mesocorticolimbic DA system is a prominent focus in research on reward processing and drug addiction, a growing literature has emerged indicating an important role for glutamate in mediating the adaptive processes underlying psychostimulant addictions. METH increases NMDAR2B synaptic currents in midbrain dopamine neurons^[34]^ and AMPAR plasticity in accumbens core contributes to incubation of METH craving^[35]^. Increasing or decreasing AMPA and NMDAR subunit trafficking to the membrane of PFC neurons during repeated exposure to psychostimulants leads to maladaptive plasticity, cognitive decline and addiction^[36]^. METH-induced selective increase in NMDAR, the increase in NMDAR current amplitude in the mPFC or increased frequency of excitatory postsynaptic currents (EPSCs) in the hippocampus contributed to the CPP^[18]^. Also, pharmacological regulation of VGLUT expression alters glutamate release and glutamate-dependent synaptic plasticity involved in drug addiction^[37]^. Our results show that VGLUT1 was increased in the PrL and dCA1 after METH-CPP expression (Figure 6), moreover silencing the pathway enhanced METH-CPP, which suggest that activity of the PrL- to-dCA1 glutamatergic pathway is critical for METH-CPP.

In conclusion, our results show that the PrL-to-dCA1 glutamatergic pathway is both necessary and sufficient to regulate METH-CPP. Moreover, we demonstrate marked changes in both the PrL and dCA1 levels of glutamate receptor subunits and VGLUT1 after METH-CPP. Future studies will examine the role of the PrL-to-dCA1 pathway in other aspects of METH conditioned learning, such as seeking induced by presenting discrete conditioned cues.

## Acknowledgments

The authors wish to thank Professor Lin Xu and Rongrong Mao for their helpful suggestion and discussion.

## Funding

National Natural Science Foundation of China (No. U2002220).

National Natural Science Foundation of China (No. 82371271).

The innovation team of stress and disorder in nervous system in Yunnan

Province (No. 202305AS350011).

## Author contributions

Conceptualization, B.J. and K.P.

Methodology, Z.D.D., S.W.J., and H.M.B.

Formal Analysis, Z.D.D., S.W.J., W.H., H.M.B., X.R., L.M.Y., G.R., B.L.P., L.Y., Z.X.W. and Z.L.

Investigation, Z.D.D., S.W.J. and W.H.

Resources, B.J.

Visualization, Z.D.D.

Funding Acquisition, B.J.

Project Administration, B.J., Z.D.D. and H.M.B.

Supervision, K.P. and B.J.

Writing – Original Draft, B.J. and Z.D.D.

Writing – Review & Editing, B.J. and K.P.

## Data and materials availability

All data are available in the main text or the supplementary materials.

## Supplementary Materials

### Materials and methods

#### Animals

Male C57BL/6 mice (8weeks old and weighing 18–25 g) were purchased from Chongqing Medical University (Chongqing, China). All mice were housed in cages (four per cage/ individually housed during photostimulation behavioral assays) under a 12-h light/dark cycle (lights on at 7 a.m.) with food and water ad libitum in a temperature (21–23°C) and humidity-controlled room. All experimental procedures were performed between 7:00 A.M. and 7:00 P.M. All experimental procedures performed in accordance with the National Institutes of Health Guide for the Care and Use of Laboratory Animals (NIH Publications No. 80-23, revised 1978). All experiments were approved by the Institutional Animal Care and Use Committee of Kunming University of Science and Technology, and the Committee on Animal Use and Protection of Yunnan province (No. LA2008305). All efforts were made to minimize the number of animals used and their suffering. Male mice were used in the current study if not indicated otherwise.

#### Viral vectors

For optogenetic experiments, we used rAAV2/9-hSyn-mCherry-WPREs-hGH pA, rAAV2/9-hSyn-hChR2(H134R)-mCherry-WPREs-hGH pA, rAAV2/9-hSyn-eNpHR3.0-mCherry-WPREs-hGH pA from HAN Bio, (Shanghai). rAAV2/9-CaMKⅡα-DIO-mCherry-WPREs-hGH pA, rAAV2/9-CaMKⅡα-DIO-hChR2(H134R)-mCherry-WPREs-hGH pA, rAAV2/9-CaMKⅡα-DIO-eNpHR3.0-mCherry-WPREs-hGH pA. For silencing of pathway, we used rAAV2/9-CaMKⅡα-DIO-TeNT-p2A-mCherry-WPREs pA, rAAV2/Retro-hSyn-Cre-EYFP-WPRE-hGH pA purchased from Brain VTA (Wuhan). For neuronal tracing and cell labelling experiments, we used rAAV2/9-(hSyn)-mCherry-WPREs-hGH pA. The titer of vectors ranges from 2.00-5.00 E+12 vg./mL.

#### Drugs and reagents

Methamphetamine (dissolved in saline) was intraperitoneally injected at a dose of 2.5 mg/kg (0.065 mL/infusion). METH (purity 84%) was provided by the Public Security Bureau of Yunnan province (Kunming, Yunnan, China) and identified using a gas chromatography-mass spectrometer. The information on primary antibody used is as follows: The c-Fos antibody (ab208942), NMDAR2B antibody (ab65783), AMPA antibody (ab109450), VGLUT1 antibody (ab227805).

#### Stereotaxic surgeries

Mice were anesthetized with intraperitoneal injection of pentobarbital (80 mg/kg), and then fixed into a stereotaxic frame. Standard surgery was performed to expose the brain surface above the PrL and dCA1. The head was shaved, and the skin was sterilized using iodine solution.

For viral injection, rAAVs were stereotaxically delivered to the following brain regions (coordinates relative to Bregma in parentheses). Coordinates used for site injection were: PrL (anterior-posterior (AP): +2.68 mm; medial-lateral (ML): +0.40 mm; dorsal-ventral (DV): −1.50 mm); dCA1 (AP: −2.30 mm; ML: ±1.80 mm; DV: −1.25 mm) according to *The Mouse Brain in Steretaxic Coordinates* by George Paxinos and Keith B.J. Franklin. The rAAVs were stereotaxically injected with a glass pipette (10 μl syringe and a 36-gauge blunt needle) connected to Nano-liter Injector (RWD Life Science Company) at a slow flow rate of 60 nl/min (unilateral volume 300 nl, injected bilaterally) to avoid potential damage of local brain tissue. After the injection we waited for at least 10 min to avoid backflow and then pull out the pipette slowly, the incision was stitched after the injection by using a surgical suture.

For optical fiber implantation, optic fiber implantation 30 min after rAAVs injections, optic fiber (200 µm in diameter, N.A. 0.37) were implanted bilaterally 1 mm above the dCA1, a microscrew was placed on the skull close to the implant site of each hemisphere to provide additional stability. Dental cement was applied to secure the optical fiber implant with a sleeve for protection. Mice remained on a heating pad until fully recovered from anesthesia and given iodophor disinfectam to the surgical wound and Penicillin sodium to fight infection (50,000U/kg, Intramuscular injection per day for 3 days. Following surgery, the mice were allowed to recover and housed at least 3 weeks before behavioral experiments, histological analyses.

For glutamatergic synaptic inactivation experiments, rAAVs were stereotaxically injected. Behavioral tests and histological analyses were conducted at 3 weeks after viral vectors injection.

#### Conditioned place preference procedure

The conditioned place preference behavior (CPP-test) apparatus consisted of two compartments (15 cm×15 cm×30 cm) with differences in visual (black or white wall) and tactile (smooth or rough floor) cues, divided by a sliding door. METH-CPP protocol was applied as our previous study, including Habituation, Pre-test, Training and Post-test sessions^[38]^. During the Habituation phase, mice were released from the middle of the conditioning apparatus and allowed to freely explore the full extent of the CPP apparatus two days for 20 min per day. Pre-test on day 3, the initial place preference was determined, mice had free access to the entire box for 20 min, the time that the mice spent in each compartment was recorded to determine the preference of experimental mice before METH administration. Generally, individual mice tended to spend a little more time in one compartment or the other during the Pre-test, thus METH was paired in the compartment in which mice trended to spend less time during the Pre-test. Training on days 4, 6, 8, 10, mice were treated with METH (2.5 mg/kg, i.p.) or saline and confined to either the white or the black compartment for 20 min. On days 5, 7, 9, 11, mice were given saline and confined to the opposite conditioning compartment for 20 min. Post-test on day 12, mice were performed without drug treatment. Time that the mice spent in each compartment was measured for 20 min and was evaluated to determine preference. After viral vectors injection or fiber implantation, the mice were individually housed for at least 3 weeks before the behavioral tests. They were handled daily by the experimenters for at least 3 days before the behavioral tests. The CPP score refers to the difference between the time lengths that each mouse spent in the drug-paired compartment in the Pre-test phase and Post-test phase. Regardless of which side the mice prefer, subtract the absolute value of 600 from the larger value of one side of the apparatus uniformly as a negative value in the calculation on the Pre-test CPP. The purpose of the rAAVs Pre-test was to exclude mice that had a exceeding preference which spent too much ≥65% (≥780 s) or ≤35% (≤420 s) of the total time (1,200 s) in one side or too few crossings (≤20 times) were eliminated before the viral injection experiment. After behavioral measure, mice were randomly assigned to either the experimental or the control group.

#### In vivo photostimulation

For photostimulation during behavioral assays, a 473 nm (blue light) or 593-nm laser (yellow light) intelligent wired photogenetic system (inper, Hangzhou) was connected to a patch cord with a pair of FC/PC connectors on each end. This patch cord was connected through a fiber optic rotary joint (which allows free rotation of the fiber; Hangzhou Newdoon Technology) with another patch cord with a FC/PC connector on one side and ceramic fiber optic cannulas on the other side (outer diameter of 1.25 mm). The ceramic fiber optic cannulas implanted in the mouse (200 μm in diameter, 0.37 NA)^[39]^ was connected to the optic patch cord using ceramic mating sleeves (Hangzhou Newdoon Technology Technology). Blue light (473 nm, 3 mW) was delivered in a train of ten, 15 ms light pulses at 20 Hz for 20 min during the Post-test in METH side. Yellow light was delivered at constant 5 mW for 20 min during the Post-test in METH side. The precise laser power was measured at the tip of the optic fiber using an Optical Power Monitor (Thorlabs).

#### Immunofluorescence staining

Mice were anesthetized and transcardially perfused with 0.9% saline for 7min followed by 4% paraformaldehyde (PFA) dissolved in 0.1 M PBS buffer (pH7.5) for 5 min. Brains were post-fixed in 4% PFA at 4°C for 4 h and then transferred to 15% for 1 day then 30% sucrose solution for 2 days. Then brains tissues were coated with embedding reagent, Tissue-Tek® O.C.T. Compound (SAKURA) and then stored in the cryoprotective buffer at −20°C. Slices were frozen sectioned into 20 μm thick.

For immunofluorescence staining, each slice was washed three times in PBS, and blocked using 5% goat serum in PBST for 1 h followed by incubation with primary antibody at 4°C overnight. After rinsing in PBS, the slices were incubated with fluorescence-conjugated secondary antibody at room temperature for 2 h. Finally, slices were coverslipped on anti-fluorescence attenuation tablets including DAPI (4’, 6-diamidino-2-phenylindole) (Solarbio). All analyses were performed blinded to the experimental conditions. In this study, the following primary and secondary antibodies were used: VGLUT1 antibody (rabbit, 1:500, Abcam, ab227805); Alexa Fluor® 488 (goat anti-rabbit, 1:500, Abcam, ab150077), images were captured using a Zeiss Axioplan 2 fluorescence microscope and AxioVision image analysis software. Images were captured using a Nikon AX fluorescence confocal microscope and NIS-Elements Viewer image analysis software (Nikon).

#### For c-Fos Immunohistochemistry

After behavioral assay, mice were sacrificed and the effect of conditioned context on the expression of c-Fos, a neuronal activity marker. In the experimental flow chart, mice were placed in the box for free movement on day 12, 90 mins after the post-cpp test of the dCA1 light stimulation was performed on the light side; The mice were anesthetized with 1.2% pentobarbital sodium reagent and treated with 0.9% saline (50 ml/mouse) and 4% paraformaldehyde reagent for cardiac perfusion (50 ml/mouse).

The whole brain tissue was taken and fixed with 4% paraformaldehyde (24 hours). After fixing, rinse the washing material with running water; The materials were dehydrated in ethanol solution of different concentrations for 2 hours, mixed with paraffin wax, and stored overnight. The coronal surface of the brain tissue was sliced for 5 μm, and the slices were spread, patch and baked (60℃, 2 h) until the wax melted, and then were dewaxed with xylene. After the hydration operation of different concentrations of alcohol (5 min), soaked in distilled water for 5 min, and to rinsed with TPBS (5 min, 3 times); During antigen repair, sections were placed in 0.01 M citric acid buffer (PH 6.0) (92 ∼ 98℃, 15 min) and rinsed with TPBS for 5 min, 3 times. Then the slices were sealed with 5% bovine serum from the same source as the second antibody (37℃, 30 min) and rinsed with TPBS for 5 min, 3 times; Standard primary antibody c-fos (1:1,000, 4℃, overnight); TPBS irrigation, 3 times; The second antibody labeled with horseradish peroxidase biotin was added (1:2,000, 37℃, 30 min) and rinsed with TPBS 3 times; color developing agent (DAB) was added for color development (lasting 10 s), rinsed with tap water to terminate color development; Hematoxylin was dyed (10 min), soaked in tap water, ethanol hydrochloride (8 s), ammonia water (15 s), and alkali returned to blue. Dehydrated in different concentrations of ethanol (5 min), transparent with xylene (10 min), sealed with neutral gum; Microscopic observation: 20×objectives optical microscope bright field observation, data collection and sorting.

#### Tissue preparation

The mice were euthanized with cervical dislocation immediately after the behavioral test was completed. By combined the map of the mouse brain, Paxinos and Franklin and the Allen Brain Reference Atlases (Adult Mouse data), the brain tissue was rapidly extracted, and then the mPFC and dCA1 were dissected bilaterally on an ice-cold plate and stored at −80°C immediately. Subsequently, all relevant proteins in different brain regions were detected with Western blotting.

#### Western blotting analysis

Protein lysates were prepared using a solubilizing solution (20 mM Tris-HCl (pH 7.4), 150 mM NaCl, 1% NP-40, 1 mM ethylene diamine tetraacetic acid (EDTA), 1 mM phenylmethanesulfonyl fluoride, 1 mM ethylene glycol bis (2-aminoethylether)-N, N’-tetraacetic acid (EGTA), 1% Triton X-100, 2.5 mM sodium pyrophosphate, 1 mM Na3VO4, 1 mM β-glycerol phosphate, and 1 mg/ml leupeptin). Protein concentration was determined using a Bio-Rad protein assay reagent (Hercules, California, USA).

Samples were separated on 12% (for β-actin, NMDAR2B, AMPAR and VGULT1 for) sodium dodecyl sulfate–polyacrylamide gel electrophoresis (SDS-PAGE) and transferred to a polyvinylidene difluoride membrane (Millipore, Billerica, Massachusetts, USA). The membrane was rinsed with PBS and nonspecific sites were blocked by incubating the membrane with blocking buffer (PBS-0.1% Tween-20 (TPBS) containing 10% nonfat milk) overnight at 4°C or for two hours at room temperature, then incubated with primary antibodies (1:1,000) followed by peroxidase-conjugated anti-mouse (1:10,000) or anti-rabbit (1:10,000) immunoglobulin G (IgG) (KPL, Inc., Gaithersburg, Maryland, USA) for one hour at room temperature. The epitope was visualized by an ECL Western blot detection kit (Millipore Corporation, Billerica, Massachusetts, USA). All analyses were performed blinded to the experimental conditions. Primary antibodies, NMDAR2B (rabbit,1:1,000, Abcam, ab65783), AMPAR (rabbit, 1:2,000, Abcam, ab109450), VGLUT1 (rabbit, 1:1,000, Abcam, ab227805). Secondary antibody, KPL Peroxidase-Labeled Antibody to Rabbit IgG (H+L) (goat anti-rabbit, 1:1,000, Seracare Life Sciences, No.5220-0336). ImageJ software was used for the densitometry analysis.

#### Imaging

Confocal fluorescence images were acquired using a Nikon AX confocal laser scanning microscope with ×10, ×20 objectives for imaging stained or autofluorescent neurons and NIS-Elements Viewer image analysis software (Nikon). fluorescence images were acquired using a Nikon TE2000-S microscope with ×10, ×20 objectives for imaging stained or auto fluorescent neurons. Coronal scans of the whole brain on PrL and dCA1 were performed using an OLYMPUS VS200 scanning microscope with ×10, ×20 objectives for imaging autofluorescent neurons and OlyVIA Viewer image analysis software (OLYMPUS). The center of viral infection was taken as the brightest fluorescent point.

#### Statistical analysis

Data were expressed as means ±SEM values. Statistical analysis was performed by using Graphpad Prism 9.0 software. Two-way ANOVA followed by a Bonferroni post hoc multiple comparison test was used to compare control and treated groups. P < 0.05 was considered to indicate a statistically significant difference.

## Supplemental Figure

**Supplemental Figure 1.**
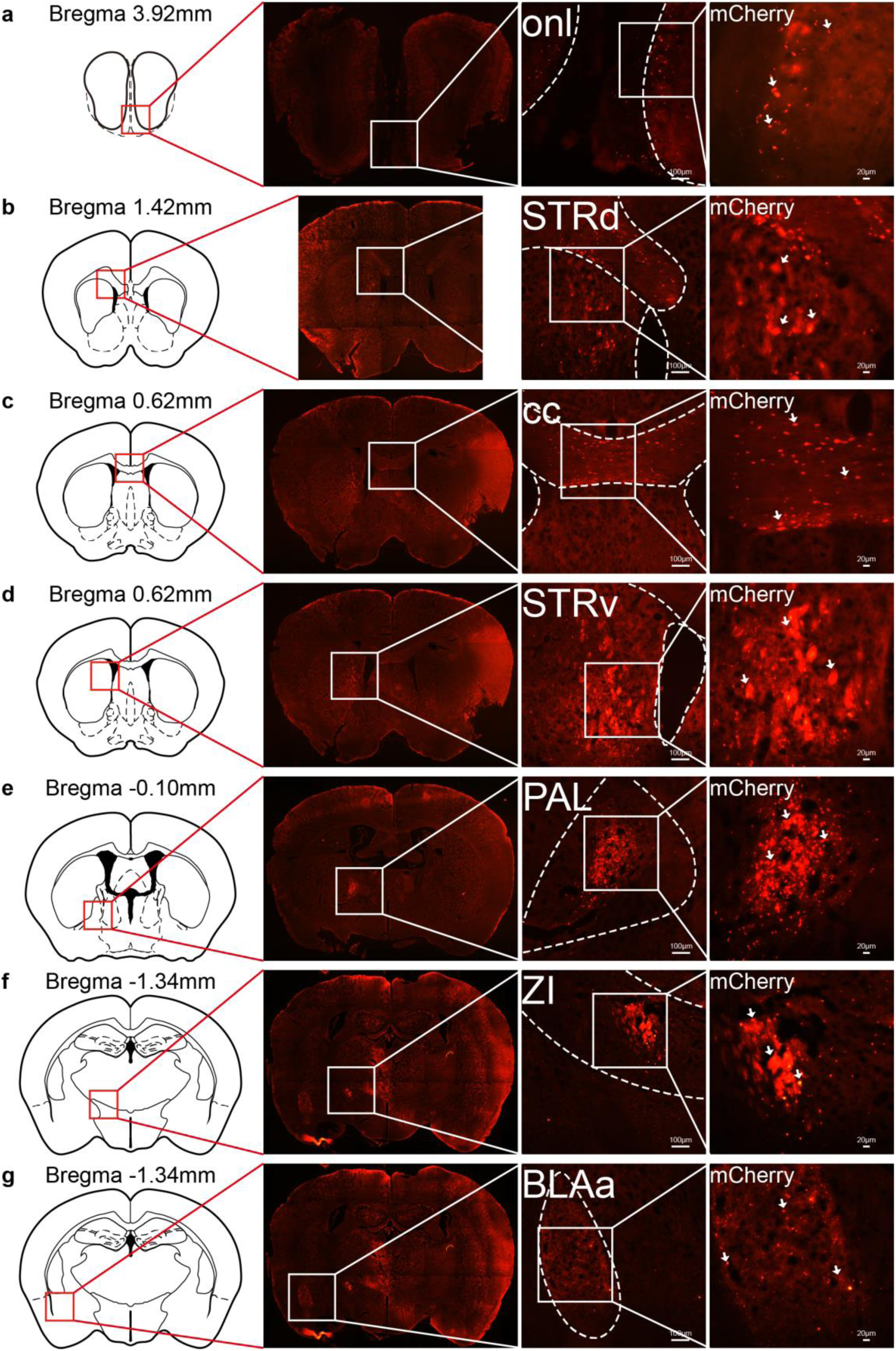
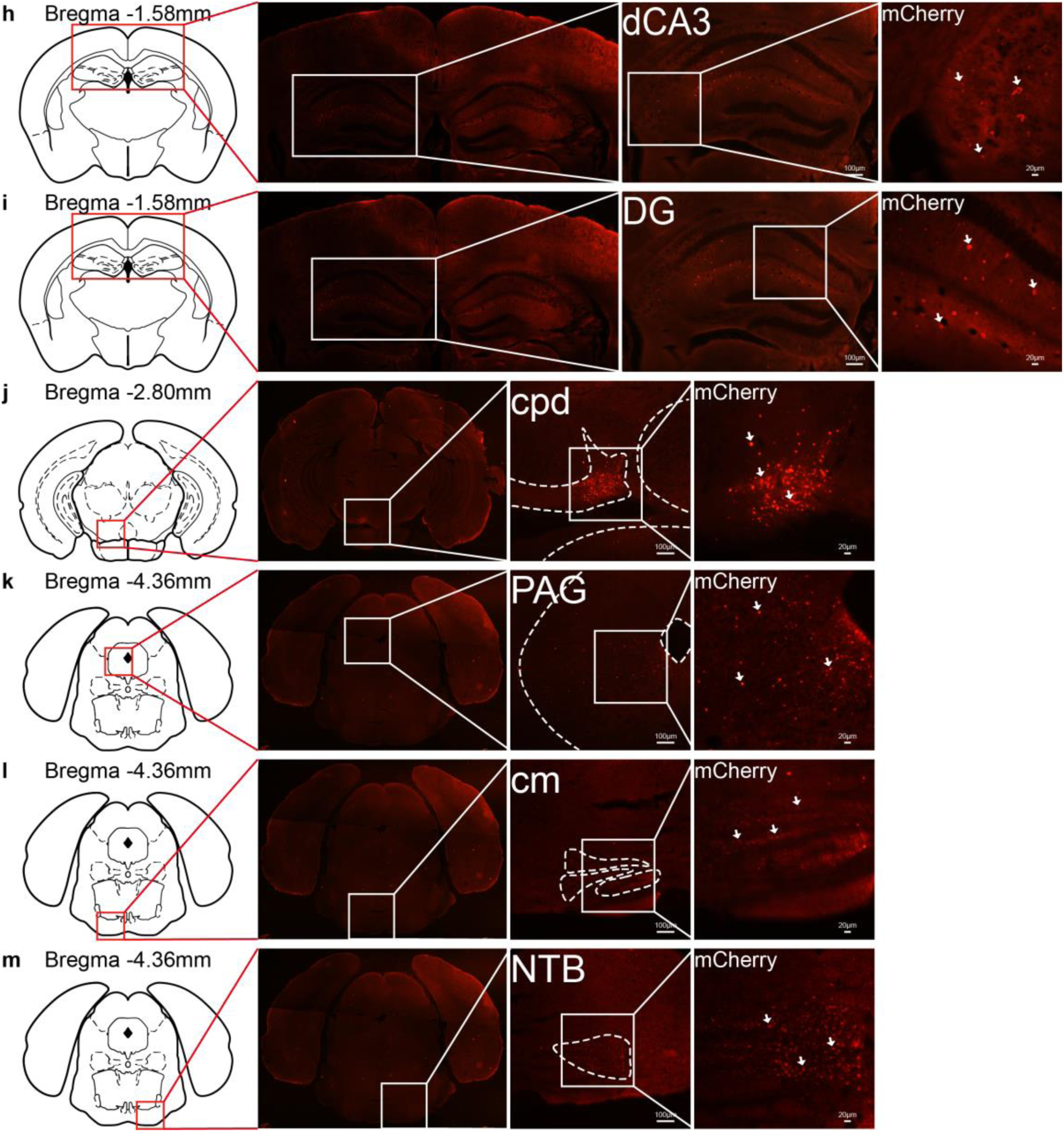
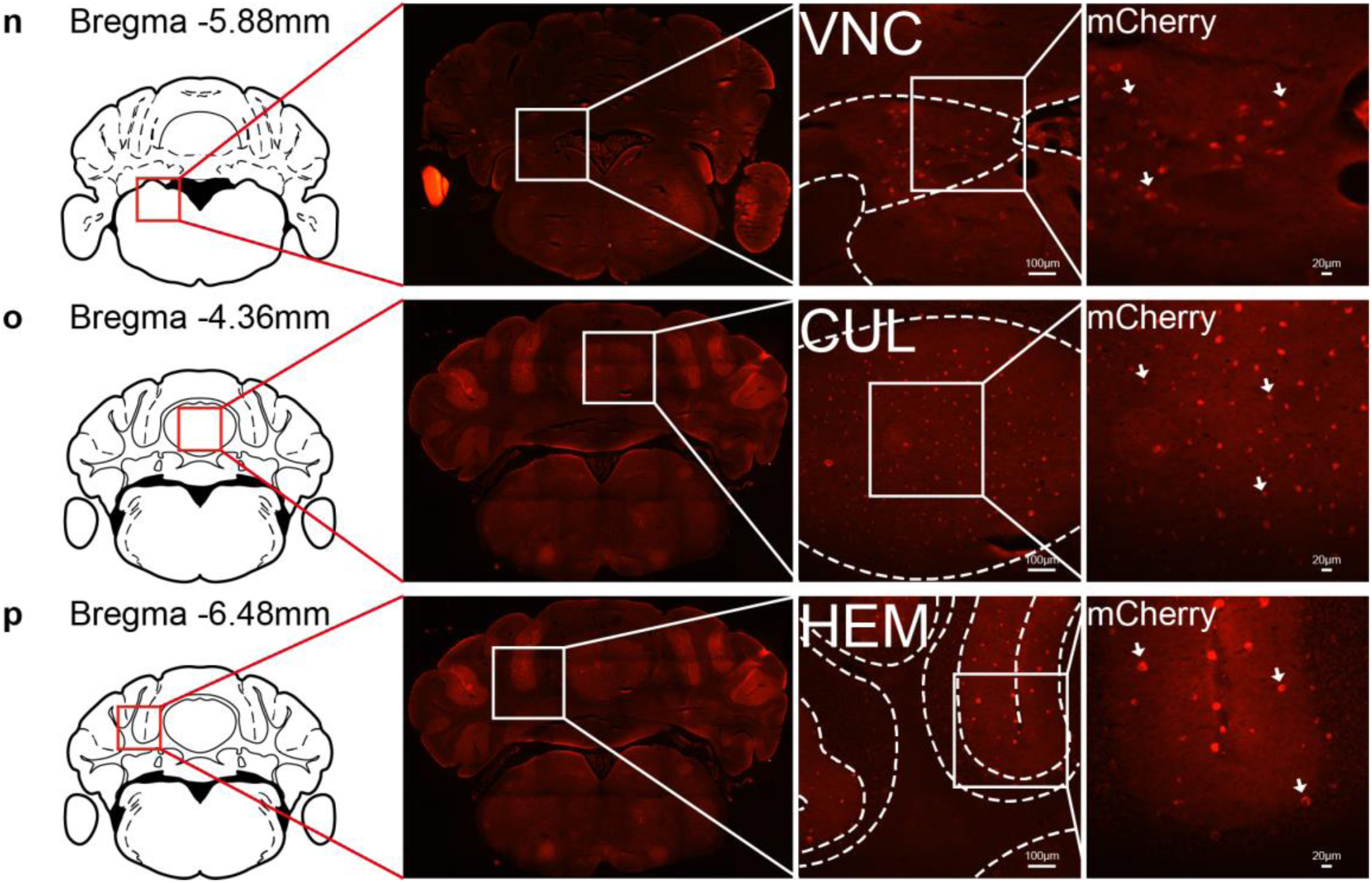
The mPFC inputs to various regions. Representative expression of the mCherry (red) in various areas of a C57BL/6 J mouse injected with rAAV2/9-hSyn-mCherry. The areas from mPFC projection. **a**, olfactory nerve layer of main olfactory bulb (onl). **b**, Striatum dorsal region (STRd). **c**, corpus callosum(cc). **d**, Striatum ventral region (STRv). **e**, Pallidum (PAL). **f**, Zona incerta (ZI). **g**, Basolateral amygdalar nucleus, anterior part (BLAa). **h**, dorsal Field CA3 (dCA3). **i**, Dentate gyrus (DG). **j**. cerebal peduncle (cpd). **k**. Periaqueductal gray (PAG). **l**. cranial nerves (cm). **m**, Nucleus of the trapezoid body (NTB). **n**, Vestibular nuclei (VNC). **o**, Culmen (CUL). **p**, Hemispheric regions (HEM). The mCherry expression in the various Regions enclosed by white boxes were shown in a higher magnification in upper right square images marked by arrow (n = 3 mice). Scale bar: 100 μm, 20 μm.

**Supplemental Figure 2.**
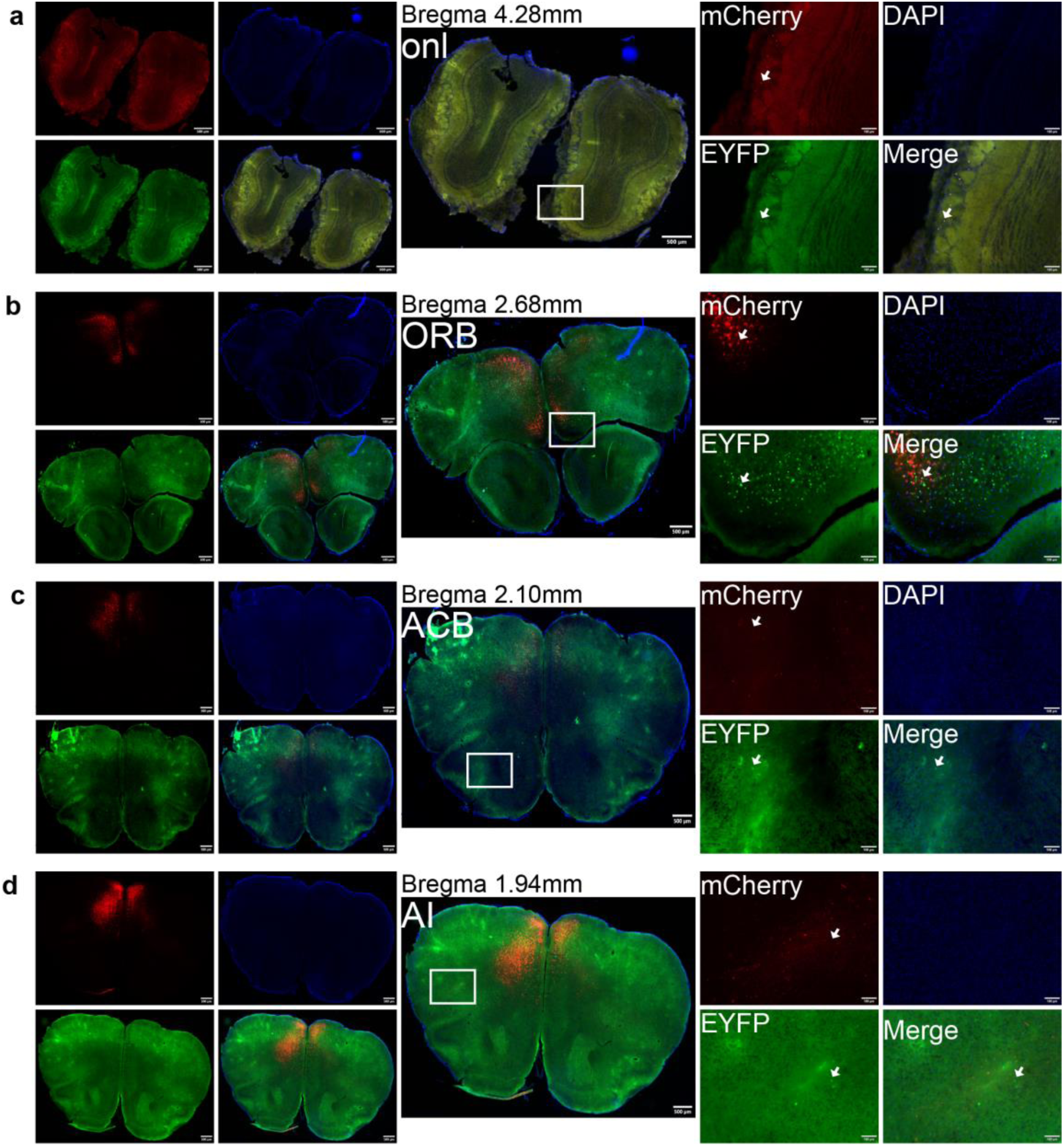
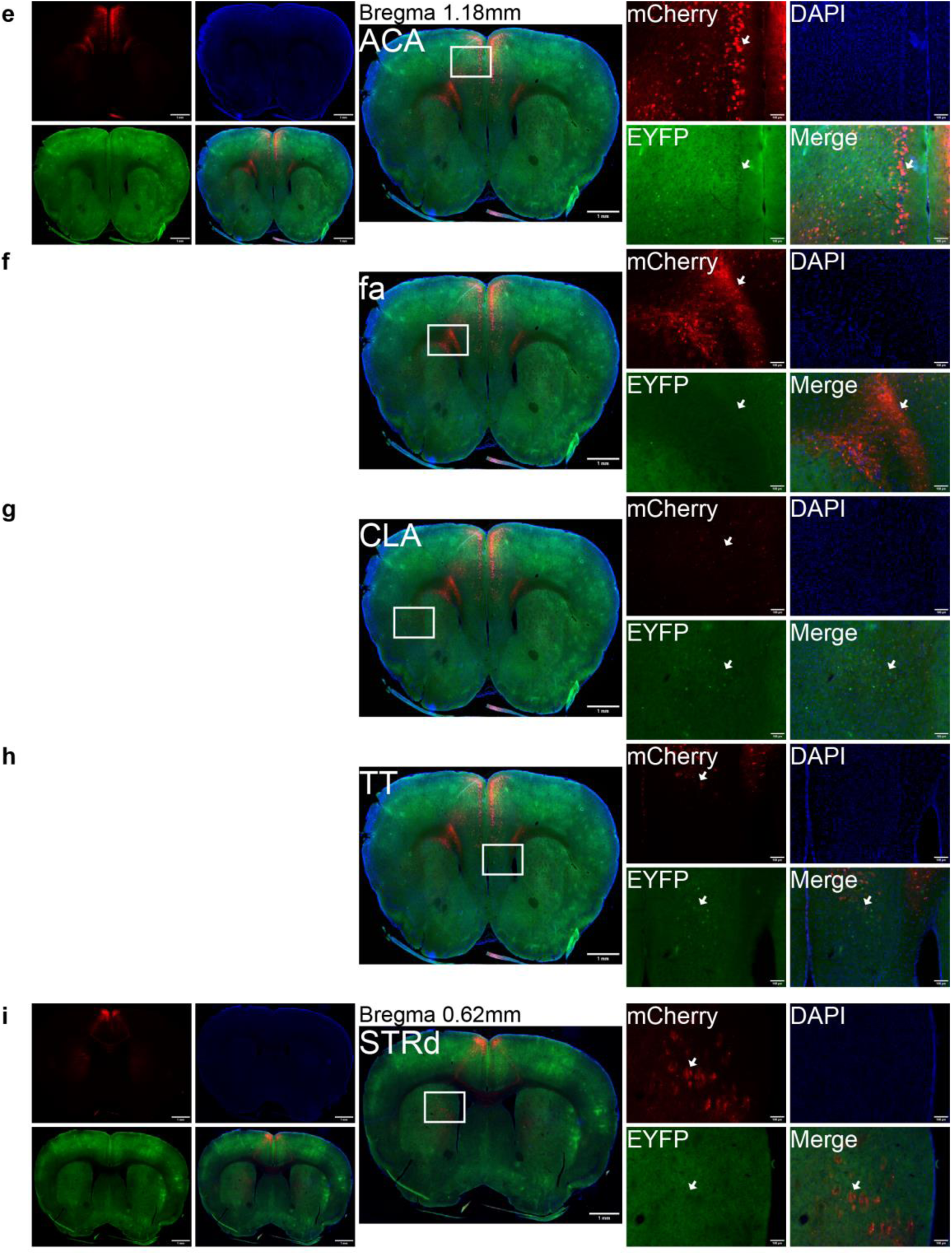
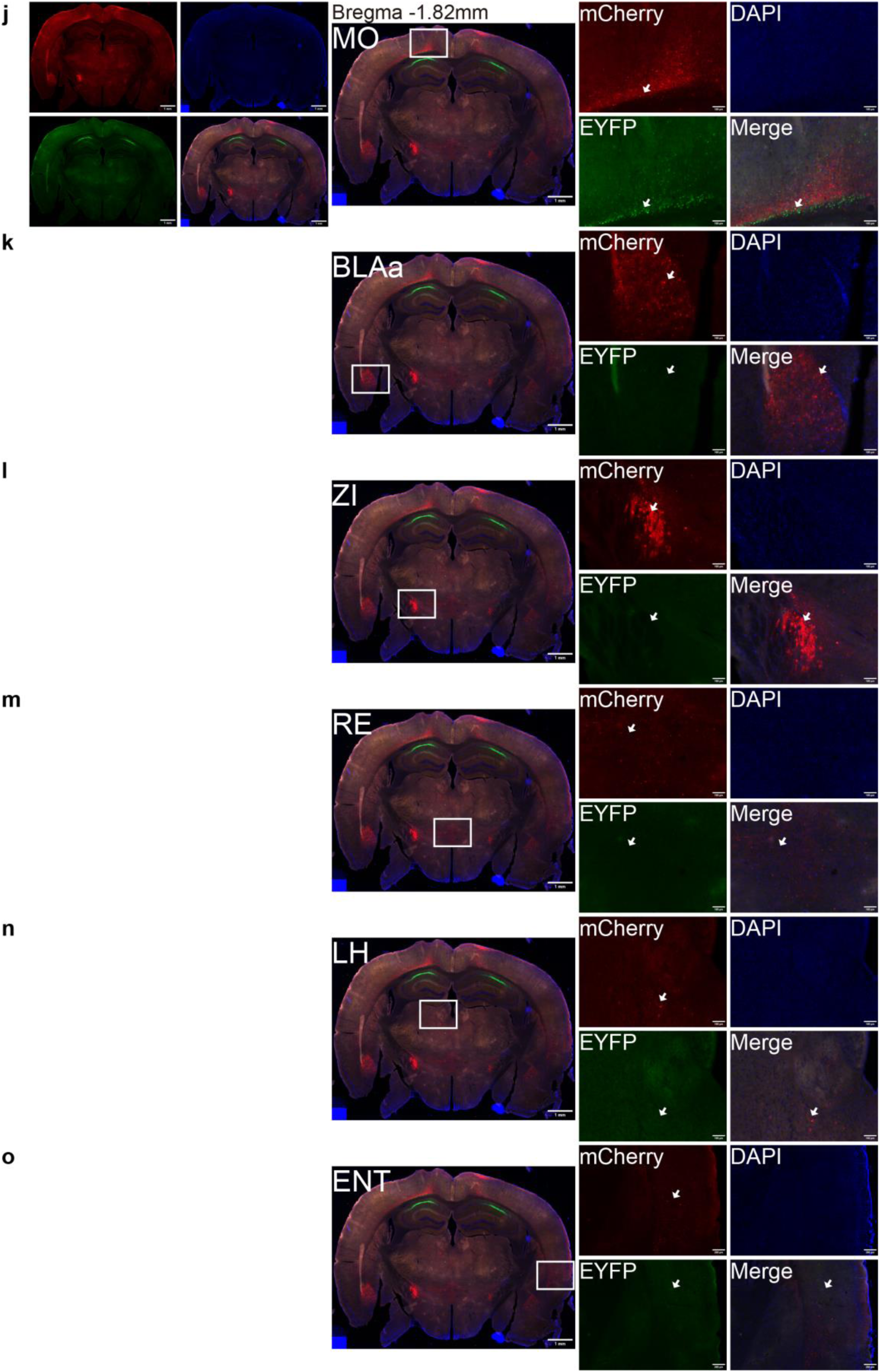
The glutamatergic PrL projects to various regions and the dCA1 inputs to various regions. Representative expression of the mCherry (red) and EYFP(Green) in various areas of a C57BL/6 J mouse injected with rAAV2/9-CaMKⅡα-DIO-mCherry in the mPFC and rAAV2/Retro-hSyn-Cre-EYFP in the dCA1. The areas from glutamatergic PrL projection and from the dCA1 input. **a**, olfactory nerve layer of main olfactory bulb (onl). **b**, Orbital area (ORB). **c**, Nucleus accumbens (ACB). **d**, Agranular insular area (AI). **e**, Anterior cingulate area (ACA). **f**, corpus callosum, anterior forceps (fa). **g**, Claustrum (CLA). **h**, Taenia tecta (TT). **i**, Striatum dorsal region (STRd). **j**, Somatomotor areas (MO). **k**, Basolateral amygdalar nucleus, anterior part (BLAa). **l**, Zona incerta (ZI). **m**, Nucleus of reuniens (RE). **n**, Lateral habenula (LH). **o**, Entorhinal area (ENT). The mCherry, EYFP and DAPI expression in the various Regions enclosed by white boxes were shown in a higher magnification in upper right square images marked by arrow (n = 3 mice). Scale bar: 500 μm, 100 μm.

